# A new pterosaur from Skye, Scotland and the early diversification of flying reptiles

**DOI:** 10.1101/2022.02.14.480264

**Authors:** Elizabeth Martin-Silverstone, David M. Unwin, Andrew R. Cuff, Emily E. Brown, Lu Allington-Jones, Paul M. Barrett

## Abstract

The Middle Jurassic was a critical time in pterosaur evolution, witnessing the inception of major morphological innovations that underpinned successive radiations by rhamphorhynchids, basal monofenestratans and pterodactyloids. Frustratingly, this interval is particularly sparsely sampled, with a record consisting almost exclusively of isolated fragmentary remains. Here, we describe new material from the Bathonian-aged Kilmaluag Formation of Skye, Scotland, which helps to close this gap. *REDACTED* (gen. et sp. nov.) is based on a three-dimensionally preserved partial skeleton, which represents the first associated Middle Jurassic pterosaur. *REDACTED* is one of the first pterosaurs to be fully digitally prepared and μCT scanning reveals multiple elements of the skeleton that remain fully embedded within the matrix, which are otherwise inaccessible. Novel anatomical features of this new Middle Jurassic pterosaur help to confirm the existence of the controversial clade Darwinoptera, greatly clarifying our understanding of Jurassic pterosaur evolution.

## 1. Introduction

The earliest known pterosaur fossils come from the Late Triassic, but the group persisted until the K-Pg extinction [1,2]. Pterosaurs are typically divided into three large groups: the basal, polyphyletic ‘rhamphorhynchoids’, which existed from the Norian; the derived pterodactyloids, which appeared in the Late Jurassic; and the basal monofenestratans that intervened between them from the Middle–Late Jurassic. Basal monofenestratans have been suggested to be a transitional, polyphyletic group leading to pterodactyloids [3,4] but the monophyly and relationships of these taxa (many referred to Wukongopteridae) are controversial [5,6].

Pterosaurs are known from every continent [2,7] and experienced two large peaks in species-richness: in the Early–’middle’ Cretaceous and latest Cretaceous [8,9]. However, their distribution is highly affected by the ‘Lagerstätten effect’ and other forms of sampling bias [8–10]. The majority of pterosaur fossils have been collected from Konservat-Lagerstätten (deposits with exceptional preservation) and it has been demonstrated that times and places of high diversity track exceptional preservation potential [9,10]. For example, the Late Jurassic pterosaur record is dominated by Konservat-Lagerstätten such as the Solnhofen Limestones of Germany and the Daohugou Beds of China, which both yield nearly complete specimens, often with associated soft tissues [1,11,12]. Both of these faunas include pterodactyloids and basal non-pterodactyloids. The younger Early–’middle’ Cretaceous is best known for areas such as the Konzentrat-Lagerstätten of the Cambridge Greensand (UK) [13], where hundreds of disarticulated and fragmentary pterodactyloid pterosaur bones have been found and several Konservat-Lagerstätten, including the Jehol Biota of China [14] and the Santana and Crato Formations of Brazil, where complete skeletons of pterodactyloids have been recovered frequently [1,11]. As a result, almost all of our knowledge of pterosaur evolutionary history is based on material from a handful of sites with restricted spatiotemporal coverage.

The Middle Jurassic pterosaur fossil record is exceptionally poor. The Stonesfield Slate (UK) provides the best example but, although abundant pterosaur material has been found, all of it is disarticulated, isolated and often fragmentary, making estimates of diversity and anatomical description difficult [15]. Recent studies have proposed that other potentially Middle Jurassic pterosaur-bearing localities, such as the Tiaojishan Formation (China), are Late Jurassic instead [16].

The scarcity of Middle Jurassic pterosaur-bearing formations and diagnosable taxa from this period are problematic when attempting to understand early pterosaur evolution. Pterosaur phylogenies suggest that Monofenestrata and Pterodactyloidea both diverged during this interval, but direct evidence for the timing of these events remains elusive. For these reasons, any new Middle Jurassic pterosaur material, especially significantly complete skeletons, are important in constraining the spatiotemporal context and evolutionary mode of the pterosaur radiation. Here, we describe a new three-dimensionally preserved partial skeleton from the Middle Jurassic (Bathonian: ~168–166 million years ago) of the Isle of Skye, Scotland, which is the first associated pterosaur discovered in Scotland and the most complete Middle Jurassic pterosaur found to date.

## 2. Material and Methods

Due to the fragility of the bones and the hardness of the surrounding matrix, traditional mechanical preparation was difficult and the specimen was partially prepared using acetic acid (see Supplementary Material). In order to visualize more details of the specimen, including elements that remain fully encased in matrix, microcomputed tomography (μCT) scanning of the specimen was performed at the University of Bristol X-Ray Tomography Facility and segmented using Avizo Lite 9.5 (see Supplementary Material for details).

The specimen was scored into a new pterosaur data matrix, consisting of 65 taxa and 137 characters, and a phylogenetic analysis was performed in TNT (see Supplementary Material for more details).

## 3. Systematic Palaeontology

PTEROSAURIA Kaup, 1834

MONOFENESTRATA Lü et al., 2010

DARWINOPTERA Andres et al., 2014

*REDACTED* gen. et sp. nov.

Holotype. NHMUK PV R37110, an associated partial skeleton consisting of several vertebrae, a sternum with cristospine, a partial right pelvis, a complete right scapulocoracoid, a partial left ulna, a proximal syncarpal, a distal syncarpal, two metacarpal IVs, several digit IV phalanx fragments, a proximal femur fragment, and numerous other long bone and unidentifiable fragments (Fig. 1).

**Figure 1:**
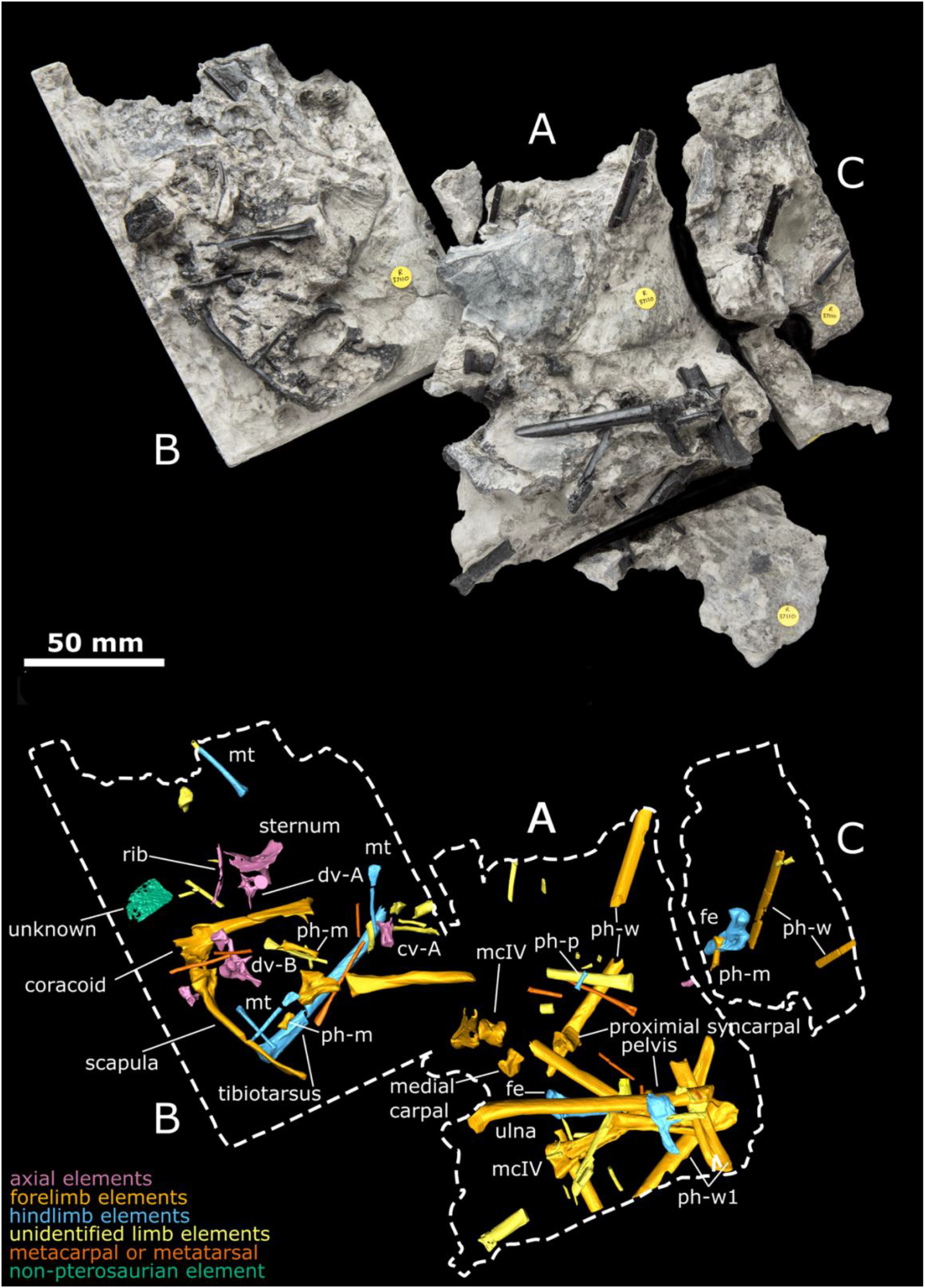
NHMUK PV R37110, REDACTED, approximately as it was found (top) and by CT reconstruction with elements (bottom). Letters indicate the blocks as discussed in the text. Abbreviations: cv/dv, cervical/dorsal vertebra; fe, femur; m, manual; mc, metacarpal; mt, metatarsal, p, pedal; ph, phalanx; w, wing.

Type locality and horizon. The holotype was found in 2006 by a team from the Natural History Museum (led by PMB), at Cladach a’Ghlinne, north of Elgol, Isle of Skye, Scotland, UK, and comes from the Bathonian-aged Kilmaluag Formation (see Supplementary Fig. 1 and [17–19]).

Etymology. [SECTION REDACTED FOR PREPRINT]

Diagnosis. Darwinopteran pterosaur (fig. 2) that differs from other darwinopterans by the presence of sternocoracoid articulations lying anterior and posterior to one another, and with a cristospine that is short and deep. *REDACTED* is further diagnosed by a single autapomorphy, the uniquely shaped end of the coracoid with an articular surface for sternal articulation.

**Figure 2:**
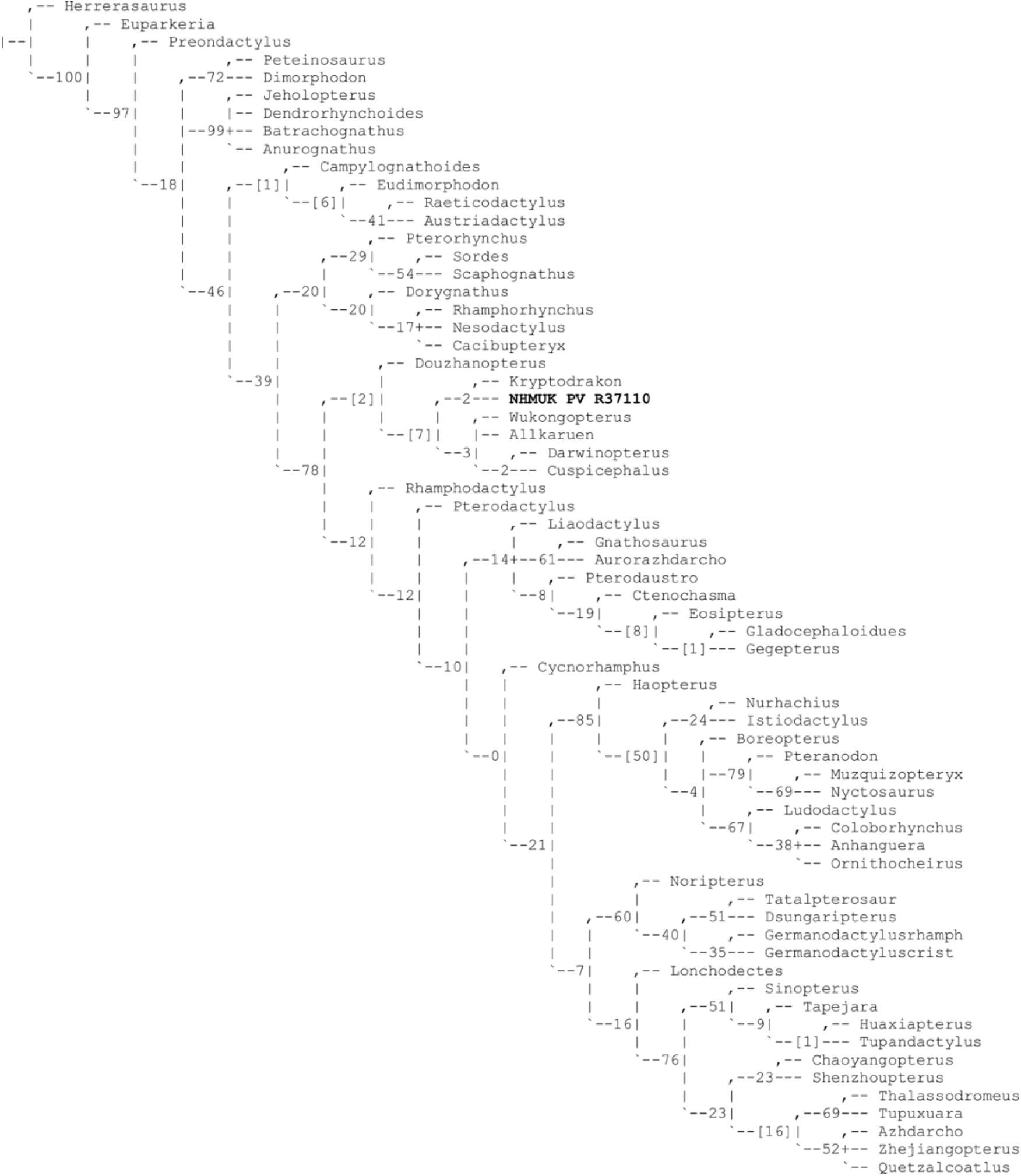
Majority rule consensus tree of 480 most parsimonious trees with bootstrap values.

## 4. Results

A more detailed description can be found in the Supplementary Material, but here we note the key anatomical features of interest. The scapulocoracoid of *REDACTED* is undistorted and complete. The scapula and coracoid are preserved as two separate yet articulated elements, but it is unclear if they were unfused or broken. The glenoid is located chiefly on the scapula, but partially on the coracoid. The scapula and coracoid are similar in length, with the coracoid being shorter (Table S1), and the distal end of the scapula is expanded and bulbous. A prominent flange is present on the distal coracoid, likely for the attachment of the *m. sternocoracoideus*. The sternum is incomplete, with just the cristospine and the most anterior portion of the sternal plate present. The cristospine is short and potential pneumatic foramina are present on the dorsal (internal) side of the base of the cristospine, where the sternal plate begins.

*REDACTED* falls out as a monofenestratan darwinopteran and sister taxon of *Kryptodrakon progenitor*. Darwinoptera is supported by five characters, including two diagnostic characters of Darwinoptera: phalanx two of pedal digit V with distinctive angular flexure at midlength, such that the distal half of the phalanx lies at 40–45°to the proximal half; and the scapula distal end is bulbous in shape. The other three features are synapomorphic reversals: ulna/tibia ratio 0.9–1.2; unguals of manus and pes similar in size; and manus digit IV (wing finger) phalanx one shorter than length of tibiotarsus.

## 5. Discussion

*REDACTED* from the Middle Jurassic of Skye, Scotland is important for several reasons. It provides new insights into key anatomical features of the shoulder girdle, manus and pes that help to unite several Jurassic pterosaurs in a single clade, Darwinoptera, which is the sister group to Pterodactyloidea. It is also the most complete undisputed Middle Jurassic pterosaur and its age and phylogenetic position have a major impact on our knowledge of pterosaur evolutionary history. So-called ‘transitional’ pterosaurs, exhibiting a mosaic of features found in the long-tailed ‘rhamphorhynchoids’ and short-tailed pterodactyloids, were first recognised with the descriptions of *Darwinopterus* [3] and *Wukongopterus* [5], which occupied varied positions in the pterosaur tree. Together with pterodactyloids, these ‘transitional’ taxa form the clade Monofenestrata [3]. These discoveries led to the proposal of modular evolution for pterosaurs, with different areas of the skeleton evolving at different times and rates [3]. *Wukongopterus* was initially regarded as a very basal pterosaur [5] and its position has been difficult to constrain [6], and several other pterosaurs have been regarded as close relatives (included in Wukongopteridae [4]). Some more recent studies have recovered a monophyletic Wukongopteridae and Darwinoptera (Wukongopteridae +*Pterorhynchus wellnhoferi)*as basal monofenestratans [20].

Our results show that Darwinoptera also includes the latest Early Jurassic/earliest Middle Jurassic pterosaur *Allkaruen koi* (previously thought to be a sister taxon to Monofenestra [21]) and the Middle–Late Jurassic *Kryptodrakon progenitor* (formerly the oldest known pterodactyloid [20]). Darwinoptera also includes taxa previously described as ‘wukongopterids’ (although they do not form a monophyletic ‘wukongopterid’ clade), including *Douzhanopterus* [4], *Cuspicephalus* [22], *Darwinopterus* [3] and *Wukongopterus* [5].

The discovery of *REDACTED* and referral of *Allkaruen koi* to Monofenestrata also results in an earlier divergence of Monofenestra and Darwinoptera than thought previously. Formerly, the earliest confirmed monofenestratans were all from the Tiaojishan Formation of China [3,4,23,24]. Although the exact age of the Tiaojishan Formation is debated, it appears to be latest Middle Jurassic (Callovian) or Late Jurassic (Oxfordian) in age, between 164–155 Ma [25–29]. Slightly younger than these are several Late Jurassic European non-pterodactyloid monofenestratans (*Cuspicephalus scarfi*, *Normannognathus wellnhoferi* [22], and the un-named ‘Painten pro-pterodactyloid’ [30]). Earlier monofenestratan records from the Bathonian Stonesfield Slate [31] are either incorrect or too fragmentary to include in phylogenetic analyses [15,20]. Our results therefore recalibrate the origin of Monofenestrata, pulling this event back to the early Middle Jurassic, approximately 8–10 million years earlier than previous estimates.

The identification of these early monofenestratans also raises questions about when and where Monofenestrata and Pterodactyloidea evolved, highlighting the patchiness of the pterosaur record at this important time in their evolution. Recovery of *Kryptodrakon progenitor* (Callovian: 162.7 Ma) as a darwinopteran, rather than the earliest pterodactyloid [20], restricts pterodactyloideans to the Late Jurassic onward, but extends the range of darwinopterans from the latest Early Jurassic to the Late Jurassic, and possibly into the Early Cretaceous by including *Wukongpterus and Kunpengopterus* within a monophyletic Darwinoptera. However, most darwinopterans are known from the latest Middle–Late Jurassic of China, leaving important gaps in the Early Jurassic. The earliest-known darwinopteran is Gondwanan [21], but our understanding of early pterosaur biogeography is limited [7].

This new tree topology also challenges our understanding of pterosaur evolution. When *Darwinopterus* was discovered, it was hailed as a transitional fossil, leading to the suggestion that pterosaurs underwent modular evolution with two phases: first, from the basal ‘rhamphorhynchoid’ pterosaurs to basal monofenestratans; with the second phase to pterodactyloids [3]. This was thought to explain the combination of features seen in *Darwinopterus*, which includes some primitive characters, such as a short wing metacarpal and a long tail, in combination with more derived features, such as a single nasoantorbital fenestra and longer skull. However, a monophyletic Darwinoptera that is the sister-taxon to Pterodactyloidea, combined with an increased number of basal monofenestratans with varying character combinations, suggests that pterosaur evolution may not have been modular, but more of a mosaic with different species evolving features at different times. Previous studies suggested that pterosaur evolution may have followed a mosaic pattern on a finer scale within anatomical units (e.g. the neurocranium and braincase) while retaining modular evolution at the level of major anatomical units (e.g. skull vs. body) [4,21]. However, we suggest that even at a gross anatomical level, modular evolution may not be tenable.

## 6. Conclusions

*REDACTED* is the first pterosaur to be named from Scotland and the most complete pterosaur to be found in the UK since Mary Anning discovered *Dimorphodon macronyx* in the early 1800s [32]. Its Middle Jurassic age, phylogenetic position and status as one of the most complete darwinopterans further elevate its significance, as it helps to shed light on when and how Monofenestrata, Darwinoptera and Pterodactyloidea evolved. Darwinoptera is shown to be a clade, not a grade, and to include pterosaurs from South America, Europe and Asia, demonstrating that it was diverse, widespread and persisted for at least 30 million years. Additional work remains to be carried out on Middle Jurassic pterosaurs, especially those from the UK. Burgeoning new vertebrate fossil finds from the late Early–Middle Jurassic of Scotland [19], especially the Isle of Skye, including dinosaurs [17,33,34], crocodylomorphs [34], ichthyosaurs [35], turtles [18], and mammals [36], suggest that this area has high potential for revealing even more critical information on Middle Jurassic faunal evolution.

## Acknowledgements

We thank S. E. Evans for introducing PMB to the locality, J. Anquetin, S. Feerick and S. Moore-Fay for assistance with fieldwork, and Scottish Nature and the John Muir Trust for permissions and access to the locality. The Natural History Museum and the Palaeontographical Society provided funding. We thank M. Day for access to the specimen, T. Davies for CT scanning, D. Sykes for an early exploratory scan and D. Martill, M. O’Sullivan, and M. Witton for discussions.

