## Supplementary material for "A new pterosaur from Skye, Scotland and the early diversification of flying reptiles"

### 1. Discovery and geological background

NHMUK PV R37110 was found on the north side of Glen Scaladal at Cladach a’Ghlinne, a small beach that forms part of the coastline of Loch Scavaig, on the Strathaird Peninsula, Isle of Skye, Scotland, UK. It was discovered during fieldwork in March/April 2006 by a party from the Natural History Museum, London, which consisted of Jérémy Anquetin, Stuart Feerick, Scott Moore-Fay and that was led by PMB.

The area is a Site of Special Scientific Interest (SSSI), which is administered by Scottish Natural Heritage (formerly Scottish Nature), and the land is owned by the John Muir Trust: fieldwork and collection permits were obtained from both organizations and are on file at the Natural History Museum, London. Collection from the cliff faces is not permitted at this locality, but permission was granted to collect from fallen blocks on the wavecut platform.

The specimen was found partially exposed as a scatter of thin-walled bones on a large boulder situated a few metres from the cliff face at the northern-most edge of the beach (close to the small headland that separates Cladach a’Ghlinne from Camasunary Bay). The specimen was collected in several pieces using hand tools and a small angle-grinder and was reconstructed in the Conservation Centre of the NHMUK (see ‘Preparation’, below).

The vertebrate-bearing horizons at this locality pertain to the Kilmaluag Formation (formerly the ‘Ostracod Limestone’) of the Great Estuarine Group, which crops out in several areas on Skye, Muck and Eigg, but reaches its maximum thickness (up to 25 m) on the Strathaird Peninsula [1–4]. Most of the known vertebrate material from the formation comes from the vicinity of Cladach a’Ghlinne (reviewed in [4–6]). The formation consists of a series of interbedded mudstones and limestones: the vertebrate remains come from the ‘Vertebrate Beds’ (or Beds 9 and 10) of Andrews [2] and are usually found in mudstone facies. Paleocene volcanic activity emplaced a number of dykes at Cladach a’Glinne, which partially metamorphosed the sediments so that although composed of mudstone they are very hard and difficult to excavate. Palaeontological, sedimentological and geochemical evidence suggests that the Kilmaluag Formation represents a series of freshwater to low-salinity environments that are often interpreted as a closed lagoonal system that fluctuated in depth and extent [2,4]. Correlations place the Kilmaluag Formation within the *retrocostatum* Zone of late Bathonian (Middle Jurassic) age [3], ~167.5–166.1 Ma ([7], updated to 2020 v. 2020/03: see https://stratigraphy.org/chart).

Cladach a'Ghlinne has yielded a rich fauna of macro- and microvertebrate remains, including hybodont sharks, osteichthyans, a ?coelacanth, caudate and albanerpetontid lissamphibians, turtles, lepidosauromorphs, choristoderes, crocodilians, non-avian dinosaurs, and a variety of mammaliaforms (reviewed in [4–6]). Isolated pterosaur teeth have been reported from the locality but remain undescribed [5].

2. **Further methods information**

   a. Preparation

The preparation of the pterosaur was challenging: the density and hardness of the matrix combined with the extreme fragility and fragmentary nature of the bones made the specimen unsuitable for mechanical preparation. Acetic acid preparation was the obvious choice for this material, but the limestone had undergone contact metamorphism, leading to uneven hardness and acid resistance.

The blocks were first rinsed with industrial methylated spirit to remove dust and cutting-tool slurry and the faces were cleaned using air-abrasive to remove carbonate deposits. The blocks were then orientated, the associated faces and breaks were marked with permanent ink, aligned using modelling clay, adhered together with Paraloid B72 in acetone [8] and allowed to cure for 4 days. The underside of the blocks were then coated in silicone rubber and supported by a cradle jacket of polyester resin and fibreglass (all resistant to acetic acid). This facilitated handling of the blocks as a single unit and prevented uncontrolled dissolution of the underside during immersion [9]. 10% acetic acid for 48 hour immersions was required to soften 1-2 mm of the most resistant areas of metamorphosed limestone. 2.5% w/v precipitated calcium phosphate was added to the bath to reduce the solution of phosphate from the fossil material [10]. This high strength acid had undesirably severe effects on softer sections of the matrix, causing undercuts and pockets to develop. This was successfully combated using local barriers: Synocryl 9123s for the bones and surrounding matrix [11] (which has since been replaced with Synocryl 9122x [12]), and microcrystalline wax for larger areas of matrix where undercuts began to develop [9].

Following each acid immersion, the blocks were rinsed in running water for 6 days to remove excess acid and calcium acetate salts [10] before mechanical removal of the softened layer with a brush and low-pressure water jet. The block was then dried in an oven for 5 hours at 50$^{o}$C to limit crystal growth. Once dry it was photographed and treated with barrier materials where necessary. The acid immersion, rinsing, oven drying, photography and masking process was repeated 29 times over the course of 12 months. Any fragments that became detached were labelled and their original location was annotated on photographs. After treatment the rubber and resin jacket was removed and a supporting mount was created using Epopast 400 epoxy laminating paste, lined with Plastazote® LD45 foam, and the specimen was housed in an acid-free cardboard box.

b. Scanning parameters and 3D model formation

An initial exploratory scan was done at the Natural History Museum Computed Tomography Facility and studied by A. Cuff. The final scans were performed at the University of Bristol in the XTM Facility of the School of Earth Sciences using a Nikon XTH 225ST X-ray tomography scanner by EM-S. with help from Tom Davies, previously of the XTM Facility. Scanning settings and parameters are as follows:

- All blocks were scanned using a 2 mm copper filter and at 8 frames per projection unless stated otherwise.
- Block A, scanned once using a reflection rotating target to a voxel size of 84.3 µm (224 kV, 357 uA, 1415 ms exposure, 3141 projections).
- Block B, scanned twice, first using a reflection target to a voxel size of 80.5 µm (224 kV, 298 uA, 2000 ms exposure, 3141 projections), second using a reflection rotating target with a 3 mm copper filter to a voxel size of 35.3 µm, to zoom into a thicker region that was difficult to visualise in the first scan (222 kV, 205 uA, 2829 ms exposure, 2850 projections).
- Block C, scanned once using a reflection target to a voxel size of 64.4 µm (222 kV, 210 uA, 1415 ms exposure, 3141 projections, 1 frame per projection)

Segmentation and visualisation of surfaces was performed using Avizo Lite 9.5 (ThermoFisher, 2018), following standard procedures [13,14]. Elements were grouped into vertebrae, forelimb, hindlimb, metatarsal/metacarpal, unknown fragments, and mystery bone groups. Colours were chosen using the Okabe and Ito Color Universal Design palette (https://jfly.uni-koeln.de/color/). Contrast in some areas was poor making segmentation slow and difficult (Figs. S1-S3). This was further complicated by pyrite which frequently infilled bones and caused numerous scanning artifacts (Fig. S1). CT scans can be downloaded from Morphosource (with downloads controlled by the NHM, Project ID 397673), while .stl files of the segmented materials of each block are included in the supplementary materials and also on Morphosource.

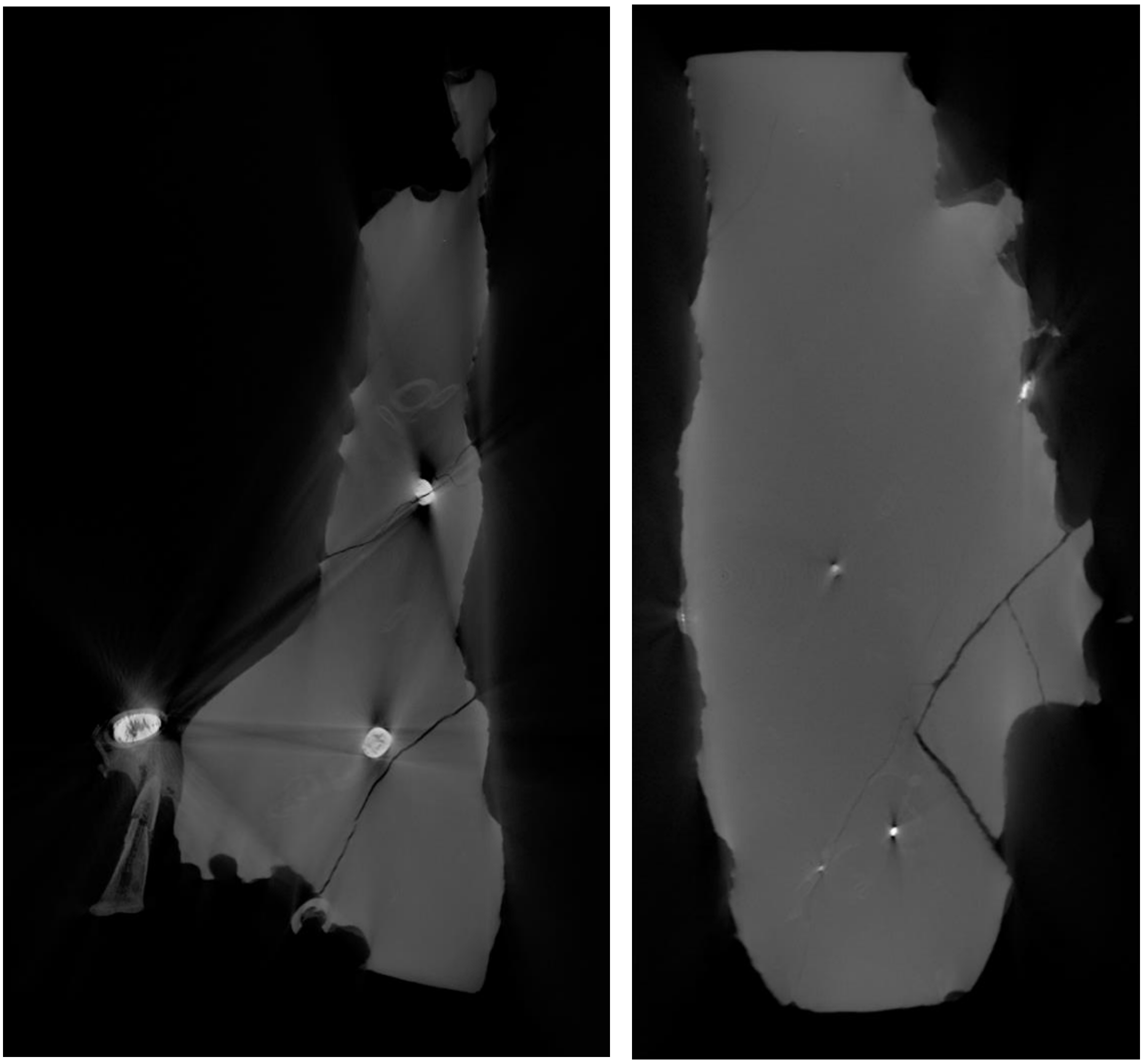

*Figure S1: Example CT slices from NHMUK PV R37110 Block A (left) and B (right) showing multiple pyrite filled bones, artifacts and poor contrast.*

c. Phylogenetic analysis

The new pterosaur data matrix was analysed in command-line TNT (Tree Analysis using New Technology: v 1.5; [15]) using the traditional (heuristic) search option with tree bisection reconnection (TBR) swapping algorithm and 10,000 replications (holding 10 trees per replication). In addition, a random seed of 2723 was used and a maximum of 1000 trees were held in memory. All characters were equally weighted and unordered. The tree search produced 480 most parsimonious trees (MPTs), from which a relatively well-resolved strict consensus tree (Supplementary Figure S2) and majority rule consensus tree (50% cut off; Figures 2, S3) were calculated. Bremer support values were calculated using the Bremer.run script available at http://phylo.wikidot.com/tntwiki#TNT_scripts (accessed: 26th October 2021). Line 177 of this script was modified to calculate bremer support values from the majority rule consensus, rather than the default strict consensus tree. Bootstrap values were calculated using the command ‘resample replications 1000 boot’.

```
Out1; mult10000=tbr hold10;
ne*
resample replications 1000 boot from 480;

GC values, 1000 replicates, cut=0 (tree 0) - Standard Bootstrap
Strict consensus of 480 trees

   ,-- Herrerasaurus
   |     ,-- Euparkeria
   |     |    ,-- Preondactylus
|--|     |    |    ,-- Peteinosaurus
   |     |    |--72--- Dimorphodon
   `--100|    |    ,-- Jeholopterus
         |    |    |-- Dendrorhynchoides
         |    |--99+-- Batrachognathus
         |    |    `-- Anurognathus
         `--97|    ,-- Eudimorphodon
              |    |-- Campylognathoides
              |    |    ,-- Raeticodactylus
              |    |--41--- Austriadactylus
              |    |              ,-- Pterorhynchus
              |    |         ,--29|    ,-- Sordes
              `--46|         |    `--54--- Scaphognathus
                   |    ,--20|    ,-- Dorygnathus
                   |    |    `--20|    ,-- Rhamphorhynchus
                   |    |         `--17+-- Nesodactylus
                   |    |              `-- Cacibupteryx
                   |    |          ,-- Douzhanopterus
                   `--39|          |     ,-- Wukongopterus
                        |    ,--[2]|     |-- Darwinopterus
                        |    |     |     |-- Kryptodrakon
                        |    |     `--[7]+-- Cuspicephalus
                        |    |           |-- Allkaruen
                        `--78|           `-- NHMUK PV R37110
                             |    ,-- Rhamphodactylus
                             |    |    ,-- Pterodactylus
                             |    |    |    ,-- Pterodaustro
                             `--12|    |    |-- Ctenochasma
                                  |    |    |-- Gladocephaloidues
                                  `--12|    |-- Gegepterus
                                       |    |-- Eosipterus
                                       |    |-- Liaodactylus
                                       |    |    ,-- Gnathosaurus
                                       `--10+--61--- Aurorazhdarcho
                                            |   ,-- Cycnorhamphus
                                            |   |         ,-- Haopterus
                                            |   |         |-- Nurhachius
                                            |   |         |-- Istiodactylus
                                            |   |    ,--85|   ,-- Boreopterus
                                            |   |    |    |   |    ,-- Pteranodon
                                            `--0|    |    |   |--80|    ,-- Muzquizopteryx
                                                |    |    `--4|    `--69--- Nyctosaurus
                                                |    |        |    ,-- Ludodactylus
                                                |    |        `--67|    ,-- Coloborhynchus
                                                |    |             `--38+-- Anhanguera
                                                `--21|                  `-- Ornithocheirus
                                                     |        ,-- Noripterus
                                                     |        |    ,-- Tatalpterosaur
                                                     |   ,--60|    |-- Dsungaripterus
                                                     |   |    `--40|    ,-- Germanodactylusrhamph
                                                     |   |         `--35--- Germanodactyluscrist
                                                     `--7|    ,-- Lonchodectes
                                                         |    |         ,-- Chaoyangopterus
                                                         |    |    ,--23--- Shenzhoupterus
                                                         `--16|    |    ,-- Huaxiapterus
                                                              |    |    |-- Sinopterus
                                                              |    |--51+-- Tupandactylus
                                                              `--76|    `-- Tapejara
                                                                   |           ,-- Thalassodromeus
                                                                   |      ,--69--- Tupuxuara
                                                                   `--[16]|    ,-- Azhdarcho
                                                                          `--52+-- Zhejiangopterus
                                                                               `-- Quetzalcoatlus
```

*Figure S2: Strict consensus tree of 480 most parsimonious trees with bootstrap values of each node.*

```
Out1; mult10000=tbr hold10;
maj*
resample replications 1000 boot from 480;

GC values, 1000 replicates, cut=0 (tree 0) - Standard Bootstrap
Majority rule consensus of 480 trees, cut 50

   ,-- Herrerasaurus
   |     ,-- Euparkeria
|--|     |    ,-- Preondactylus
   |     |    |         ,-- Peteinosaurus
   `--100|    |    ,--72--- Dimorphodon
         |    |    |    ,-- Jeholopterus
         `--97|    |    |-- Dendrorhynchoides
              |    |--99+-- Batrachognathus
              |    |    `-- Anurognathus
              |    |          ,-- Campylognathoides
              `--18|    ,--[1]|     ,-- Eudimorphodon
                   |    |     `--[6]|    ,-- Raeticodactylus
                   |    |           `--41--- Austriadactylus
                   |    |              ,-- Pterorhynchus
                   |    |         ,--29|    ,-- Sordes
                   |    |         |    `--54--- Scaphognathus
                   `--46|    ,--20|    ,-- Dorygnathus
                        |    |    `--20|    ,-- Rhamphorhynchus
                        |    |         `--17+-- Nesodactylus
                        |    |              `-- Cacibupteryx
                        |    |          ,-- Douzhanopterus
                        `--39|          |         ,-- Kryptodrakon
                             |    ,--[2]|     ,--2--- NHMUK PV R37110
                             |    |     |     |   ,-- Wukongopterus
                             |    |     `--[7]|   |-- Allkaruen
                             |    |           `--3|   ,-- Darwinopterus
                             `--78|               `--2--- Cuspicephalus
                                  |    ,-- Rhamphodactylus
                                  |    |    ,-- Pterodactylus
                                  |    |    |         ,-- Liaodactylus
                                  `--12|    |         |    ,-- Gnathosaurus
                                       |    |    ,--14+--61--- Aurorazhdarcho
                                       |    |    |    |   ,-- Pterodaustro
                                       `--12|    |    `--8|    ,-- Ctenochasma
                                            |    |        `--19|     ,-- Eosipterus
                                            |    |             `--[8]|     ,-- Gladocephaloidues
                                            |    |                   `--[1]--- Gegepterus
                                            `--10|   ,-- Cycnorhamphus
                                                 |   |         ,-- Haopterus
                                                 |   |         |           ,-- Nurhachius
                                                 |   |    ,--85|      ,--24--- Istiodactylus
                                                 |   |    |    |      |   ,-- Boreopterus
                                                 `--0|    |    `--[50]|   |    ,-- Pteranodon
                                                     |    |           |   |--79|    ,-- Muzquizopteryx
                                                     |    |           `--4|    `--69--- Nyctosaurus
                                                     |    |               |    ,-- Ludodactylus
                                                     |    |               `--67|    ,-- Coloborhynchus
                                                     `--21|                    `--38+-- Anhanguera
                                                          |                         `-- Ornithocheirus
                                                          |        ,-- Noripterus
                                                          |        |         ,-- Tatalpterosaur
                                                          |   ,--60|    ,--51--- Dsungaripterus
                                                          |   |    `--40|    ,-- Germanodactylusrhamph
                                                          |   |         `--35--- Germanodactyluscrist
                                                          `--7|    ,-- Lonchodectes
                                                              |    |         ,-- Sinopterus
                                                              |    |    ,--51|   ,-- Tapejara
                                                              `--16|    |    `--9|     ,-- Huaxiapterus
                                                                   |    |        `--[1]--- Tupandactylus
                                                                   `--76|         ,-- Chaoyangopterus
                                                                        |    ,--23--- Shenzhoupterus
                                                                        |    |           ,-- Thalassodromeus
                                                                        `--23|      ,--69--- Tupuxuara
                                                                             `--[16]|    ,-- Azhdarcho
                                                                                    `--52+-- Zhejiangopterus
                                                                                         `-- Quetzalcoatlus
```

*Figure S3: Majority rule consensus tree of 480 most parsimonious trees with bootstrap values.*

### 3. Further description

Here we provide a more detailed anatomical description of the elements known for *REDACTED.* A skeletal drawing is also provided to show which aspects of the skeleton are present and to what degree they are preserved (Fig. S4). The description will be listed by which elements are found in each block, in order to make it clear where each element lies. Elements are mostly disarticulated but associated, with a number of overlying elements. In many cases, the elements go into the matrix or are fully enclosed within the matrix, meaning that CT scanning was necessary to identify and see many elements. In the description of each elements, details of preservation and position will also be included. Measurements for significantly preserved elements found in Table S1.

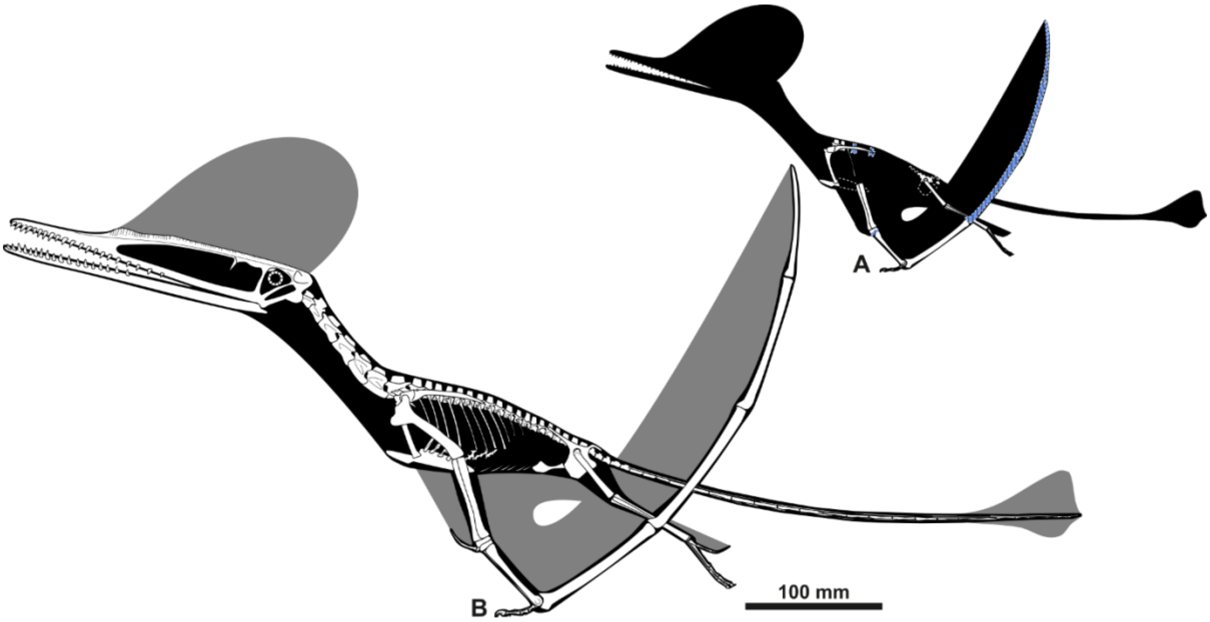


*Figure S4: Skeletal drawing of* REDACTED *showing which elements are present (A) and a hypothetical whole skeleton (B) based on closely related taxa. Blue hatched elements in A indicate elements that are likely present but incomplete and/or poorly preserved. Image by Mark Witton.*

a. Block A (Figs. 1, S5)

Pelvis (R) – postacetabular process, some ischium and ilium

Left Ulna (L) – Mostly preserved on the surface, with (which end) preserved.

Proximal Syncarpal (L) – The left proximal syncarpal is complete, preserved entirely within the matrix and not visible without CT scanning. It is completely fused into a single element. It is not articulated, but preserved nearby the medial carpal below.

Medial Carpal – The medial carpal is also complete and preserved entirely within the matrix.

Left Metacarpal IV – The left MCIV is complete, though slightly crushed in the diaphysis. It is entirely preserved within the matrix. It is reasonably elongate, with a length:width ratio of 9:1, and a total length of 57 mm (Table S1).

Right Metacarpal IV – The right MCIV is incomplete, preserving only approximately the distal third, including the distal articular end where MCIV articulates with WP1. The cortical thickness in the diaphysis where it is broken is 0.93 mm in the thinnest section and 1.82 mm in the thickest. This end is visible on the surface.

Metacarpal/metatarsal of digit III – incomplete metacarpal or metatarsal preserving one articular end (which one?)

Wing phalanx (WP) 1 – Proximal end of left(?) WP1

Wing phalanx 1 – Distal end of WP1 (maybe left, maybe same as proximal above)

Wing phalanx fragments – several fragments of wing phalanx, showing characteristic cross-sectional shape and thickness.

Femur (R)– significant portion of diaphysis and the distal articular end of the right femur.

Pedal phalanx – complete.

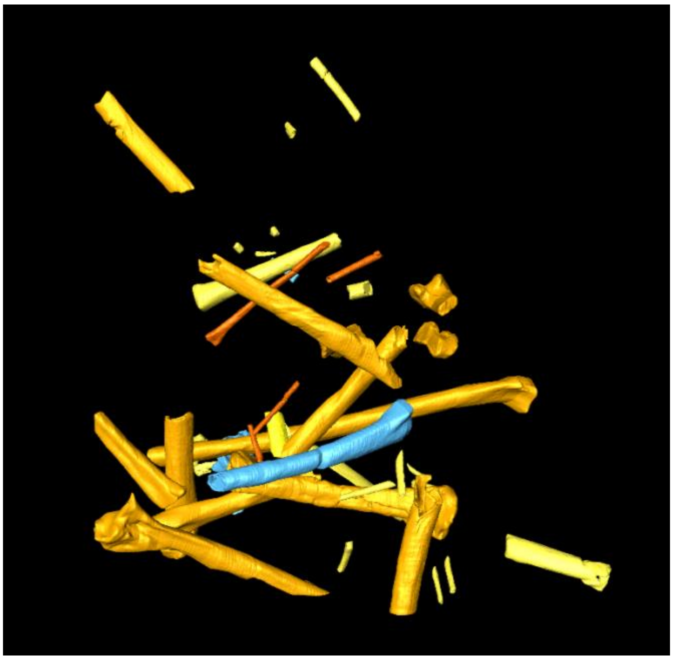

*Figure S5: underside of CT rendered Block A, NHMUK PV R37110.*

b. Block B (Figs. 1, S6-8)

Sternum – with double facets, one above the other. Short, robust cristospine, deep sternum. Preserved entirely within the matrix.

Rib – A partial, poorly preserved early rib, but preserving the articular end.

Dorsal vertebra (DV) A – Preserved entirely within the matrix, next to the coracoid. Well preserved centrum, neural canal, zygapophyses, and a small neural spine.

Dorsal vertebra B – Centrum visible on the surface of the block, present between the scapula and coracoid. Well preserved centrum, neural canal, zygapophyses, and a larger neural spine than DVA.

Dorsal vertebra C – Only visible in the higher resolution scans (Figs. S7-S8), fully within the matrix. Significantly curved/pinched centrum in ventral view.

Dorsal vertebra D – Only visible in higher resolution scans (Figs. S7-S8), fully within the matrix. Well preserved centrum, neural canal, zygapophyses, and neural spine.

Caudal vertebra A – similar morphototype to cervical vertebrae, likely one of the anterior caudals 1-3.

Vert A – A very partial poorly preserved vertebra in the apex of the scapulocoracoid. It includes part of the centrum and potentially part of the neural spine.

Vert B - Preserved at the top of the scapula. Poorly preserved.

Scapulocoracoid (R) – Complete, preserved on the surface of the block. Both elements are complete, and articulated, but separate, and it is unclear if they were unfused or broken. The glenoid is preserved mainly on the scapula, but also partially on the coracoid.

i. Scapula – Distal end is expanded and bulbous

ii. Coracoid – prominent flange for attachment of *m. sternocoracoideus* present on the distal (?) end.

Ulna/radius – not very well preserved, unsure if it is an ulna or radius, mostly just diaphysis.

Ulna (?) proximal end - very poorly preserved, next to tibiotarsus and possible ulna/radius.

Manual phalanx (penultimate) - elongated, both articulation points present.

Manual phalanx III (basal) – well preserved, complete.

Manual ungual – preserved on the underside of the block (not visible in Fig. 1).

Tibiotarsus (L) – Partial tibia with distal articulation preserved and visible on underside, with distance of diaphysis present.

Metatarsals - Four partial metatarsals preserved with the articulation end, and two being very robust looking.

Fragments of metacarpal or metatarsal diaphyses.

Unknown/non-pterosaurian bone – On the surface is a rectangular/square-ish bone that is of unknown origin. It is unlike any pterosaur bone, being porous and unusual. We suspect it is something like a crocodilian scute or a piece of turtle plastron or carapace.

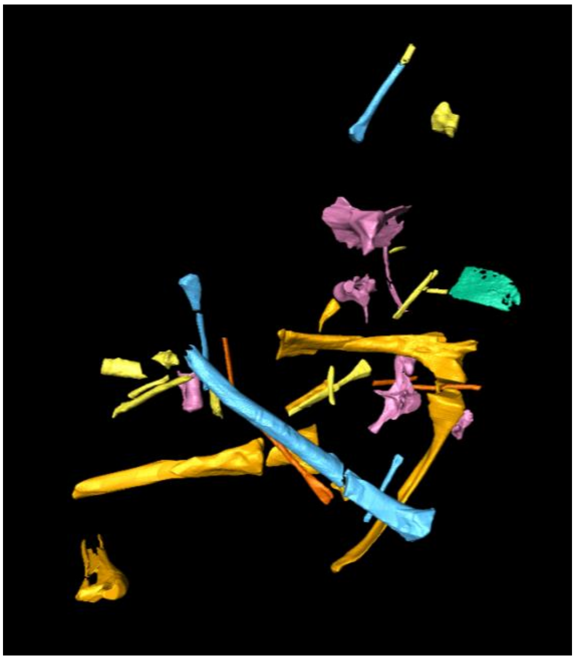

*Figure S6: underside of CT rendered Block B, NHMUK PV R37110.*

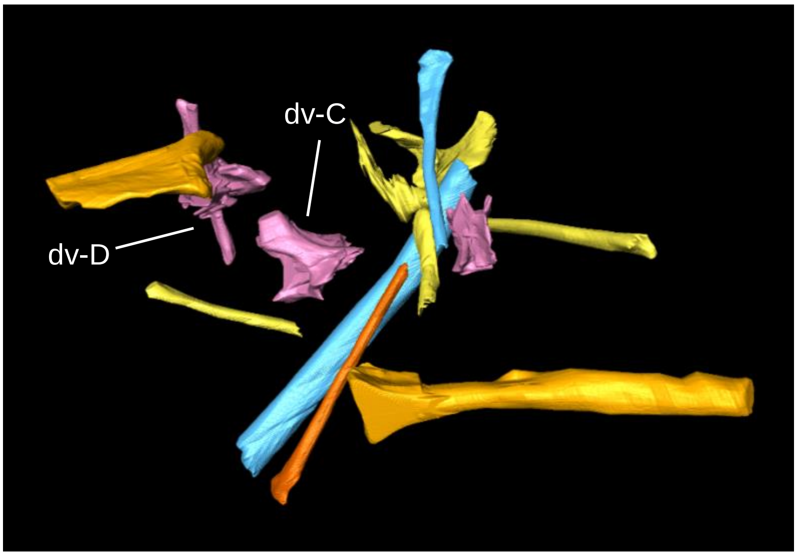


*Figure S7: High resolution of CT rendered Block B. Note vertebrae not present in the lower resolution scan seen in Figs. 1, S6.*

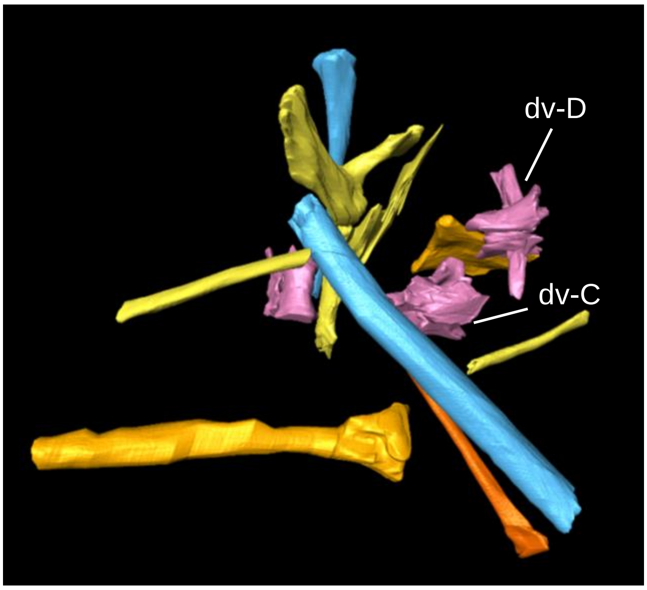


*Figure S8: Underside of high resolution CT rendered Block B (Fig. S6). Note vertebrae not visible in lower resolution scans (Figs. 1, S6).*

c. Block C (Figs. 1, S9)

Caudal vertebra – fragment near the femur.

Femur (R) proximal – proximal end and head of the right femur, at an angle of 126°.

Manual Phalanx – highly curved

Wing phalanx fragments - fragments from either wing phalanx 3 or 4.

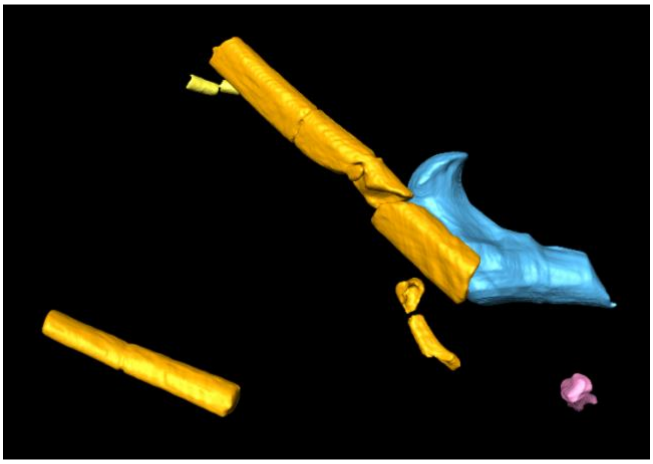

*Figure S9: Underside of CT rendered Block C (Fig. 1).*

d. Isolated/minor blocks

Distal end of coracoid (L) – preserving the articular end and the flange for *m. sternocoracoideus* attachment (Fig. S10).

Fragments of wing phalanx and other wing bones.

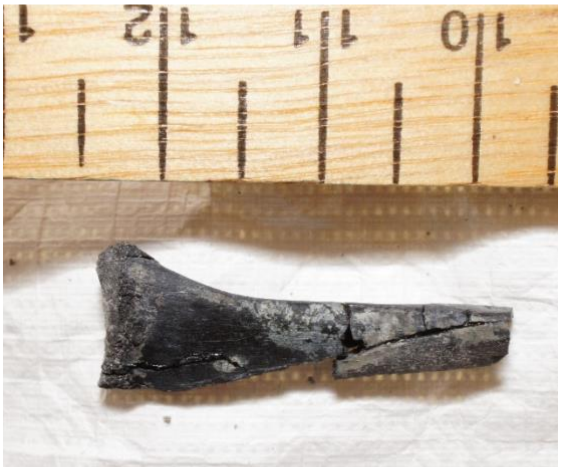

Figure S10: Isolated distal end of the left coracoid from NHMUK PV R37110.

Table S1: Measurements of selected elements. Ct = cortical thickness, CL = centrum length. *indicates incomplete bone, maximum preserved length measured. † indicates average between two different measurements (greatest and smallest). a indicates maximum length preserved between the ends of two zygapophyses in a vertebra. b indicates maximum height preserved from the base of the centrum to the top of the neural spine in a vertebra.

| | Length (mm) | Diameter, mid-diaphysis if possible (mm) | Additional measurements (mm) |
|---|---|---|---|
| **Block A** | | | |
| Ulna (L) | 102 | 5 | Ct = 0.64† |
| WP1 (L) prox | 60* | 7 | Ct = 0.88† |
| WP1 dist | 40* | 7 | Ct = 0.90† |
| WMC (L) | 57 | 6 x 3 | Ct =0.89 |
| WMC (R) dist | - | 7 x 3 | Ct = 0.93 x 1.82 |
| Femur (R) dist | 53* | 4 | Ct = 0.72 |
| Pedal phalanx | 8 | 1.4 | |
| **Block B** | | | |
| Scapula (R) | 57 | 6 | |
| Coracoid (R) | 51 | 4 | |
| Tibiotarsus (L) | 65* | 4 | Ct = 0.76† |

| | | | |
|---|---|---|---|
| Manual ph | 13 | 1.5 | |
| DV-A | 13[a] | 11[b] | CL = 7 |
| DV-B | 20[a] | 12[b] | CL = 8 |
| DV-C | | 14[b] | CL= 11 |
| DV-D | 20[a] | 12[b] | CL= 9 |
| **Block C** | | | |
| Manual ph | 11 | 1.6 | |
| Femur (R) prox | | 5 | Ct = 0.64 |
